# Identification of PAM Requirements for the *Vibrio cholerae* type I-E CRISPR-Cas System

**DOI:** 10.64898/2025.12.24.696430

**Authors:** Anne M. Stringer, Joseph T. Wade

## Abstract

CRISPR-Cas systems are prokaryotic adaptive immune systems that use RNA-guided protein complexes to target invading nucleic acid. A surveillance complex consisting of protein and a CRISPR-RNA (crRNA) binds target nucleic acid via base-pairing interactions, typically leading to processing of the target nucleic acid by a nuclease. CRISPR-Cas systems are classified based on their mechanism of action, with type I systems being the most prevalent in nature. Type I CRISPR-Cas systems target DNA, and require extensive complementarity between the crRNA and the target DNA. Moreover, type I systems require the presence of a “Protospacer Adjacent Motif” (PAM) sequence in the target DNA immediately adjacent to the expected region of base-pairing with the crRNA. Classical biotypes of the bacterial pathogen *Vibrio cholerae* have active type I-E CRISPR-Cas systems. While the optimal PAM sequence for this CRISPR-Cas system is known to be AAY, the activity of other sequences as possible PAMs has not been determined. Here, we quantify the effectiveness of all possible trinucleotide sequences in the PAM position for the *V. cholerae* type I-E CRISPR-Cas system. Our data indicate a hierarchy of PAM efficacy, with 15 of the 64 trinucleotide sequences functioning as a PAM.

## INTRODUCTION

CRISPR-Cas systems are prokaryotic adaptive immune systems that protect against invading nucleic acid such as bacteriophage and plasmids (Nussenzweig and Marraffini, 2020). The hallmark of CRISPR-Cas systems is an RNA-guided surveillance complex. The RNA component, referred to as the CRISPR-RNA (crRNA), provides sequence-specificity for target nucleic acid, which can be DNA or RNA depending on the specific CRISPR-Cas system. When a crRNA is base-paired with a target DNA/RNA in the context of the surveillance complex, the target DNA/RNA is typically cleaved by a nuclease activity of the surveillance complex or an associated protein, in a process referred to as “interference”. Thus, CRISPR-Cas immunity can be predicted by comparing crRNA sequences to those of potential target DNAs.

crRNAs are transcribed from CRISPR arrays, alternating DNA units of “repeats” and “spacers”. Repeat sequences are identical; spacer sequences are the same or similar lengths to one another but differ in their sequence. Individual crRNAs are processed out of CRISPR array transcripts by nucleases that cut within the repeat sequences. Thus, each crRNA contains a single, complete spacer sequence; it is this spacer sequence that will base-pair with the target DNA in the context of the surveillance complex. The target DNA/RNA site is referred to as the “protospacer”. Most CRISPR-Cas systems also require a “Protospacer Adjacent Motif” (PAM) in the target sequence to facilitate base-pairing with the crRNA; the PAM is usually located immediately adjacent to, or close to the protospacer (Gleditzsch *et al*., 2019).

CRISPR-Cas systems are adaptive, meaning that new spacers and repeats can be added to a CRISPR array when an organism encounters an invading nucleic acid molecule. The process of adding new repeats and spacers to a CRISPR array is referred to as “adaptation”, and requires the Cas1 and Cas2 proteins. For some CRISPR-Cas systems, the process of adaptation can be coupled to interference, in a process known as “primed adaptation”.

There are two classes of CRISPR-Cas systems: Class I systems, in which the surveillance complex consists of multiple proteins, and Class II systems, in which the surveillance complex is a single protein with inherent nuclease activity (Makarova *et al*., 2020). These Classes are further subdivided into types. The predominant type of CRISPR-Cas system encoded in sequenced prokaryotic genomes is the type I system, which targets DNA, and is characterized by the presence of an accessory nuclease protein, Cas3. Type I systems are further subdivided into subclasses A-G based on the protein composition of the surveillance complex. Arguably the best-studied type I CRISPR-Cas system is the type I-E system, which comprises a multi-protein surveillance complex referred to as “Cascade” (Brouns *et al*., 2008). When a crRNA base-pairs with protospacer DNA in the context of the Cascade complex, the Cas3 nuclease is recruited to degrade the target DNA (Xiao *et al*., 2018). Type I-E systems use a 3 bp PAM that is immediately adjacent to the protospacer. High-resolution structures of Cascade bound to protospacer DNA have been determined for the complexes from *Escherichia coli* (Hayes *et al*., 2016) and *Thermobifida fusca* (Xiao *et al*., 2017). In both cases, the DNA minor groove at the PAM interacts with the Cse1 protein of Cascade. While the precise molecular details of PAM binding differ for the two complexes, both use a conserved glutamine to form a “wedge” that separates the two DNA strands to facilitate crRNA base-pairing with the target DNA strand (Hayes *et al*., 2016; Xiao *et al*., 2017). The optimal PAM for the *E. coli* type I-E system has been determined to be 5’-AAG-3’ / 5’-RTT-3’, where AAG is on the non-target strand that does not base-pair with the crRNA. Henceforth, we will refer to PAM sequences on the non-target strand only. The PAM is also recognized by the *E. coli* Cas1-Cas2 complex (Nuñez *et al*., 2014; Wang *et al*., 2015), and hence is required for efficient adaptation (Swarts *et al*., 2012; Datsenko *et al*., 2012; Savitskaya *et al*., 2013; Fineran *et al*., 2014; Shmakov *et al*., 2014; Strotskaya *et al*., 2017; Stringer *et al*., 2020b).

Type I-E CRISPR-Cas systems are found throughout the classical biotypes of *Vibrio cholerae* and have been shown to be active against lytic bacteriophage (Box *et al*., 2016) and conjugated plasmids (Stringer *et al*., 2020a; Bourgeois *et al*., 2020). Sequence analysis of *V. cholerae* spacers has identified hundreds of likely protospacers in bacteriophage, prophage, and plasmids (Box *et al*., 2016; Bourgeois *et al*., 2020). These protospacers tend to be associated with an AAY PAM. Mutations in an AAY PAM associated with a plasmid-borne protospacer reduced the ability of the *V. cholerae* CRISPR-Cas system to prevent plasmid conjugation (Bourgeois *et al*., 2020). Thus, there is strong evidence from independent studies that AAY is the optimal PAM. However, there is limited information on the ability of other sequences to function as PAMs. The *E. coli* type I-E CRISPR-Cas system retains partial function with several non-optimal PAM sequences (Westra *et al*., 2013; Fineran *et al*., 2014; Xue *et al*., 2015; Leenay *et al*., 2016), suggesting that sequences other than AAY could function as PAMs in *V. cholerae*, albeit less efficiently than AAY. Knowing the full set of possible PAM sequences has important implications. First, suboptimal PAMs can promote primed adaptation (Datsenko *et al*., 2012; Fineran *et al*., 2014; Xue *et al*., 2015). Second, suboptimal PAMs may be useful for tuning the activity of type I-E CRISPR-based regulatory systems (Caliando and Voigt, 2015). Third, a list of sequences that can function as PAMs would facilitate analysis of putative protospacers. Here, we use a library of plasmid-borne protospacers to quantify the effectiveness of all trinucleotide sequences as PAMs for the *V. cholerae* A50 CRISPR-Cas system.

## RESULTS AND DISCUSSION

We previously showed that the activity of the *V. cholerae* CRISPR-Cas system towards different protospacer sequences can be determined by conjugating a library of plasmid-borne protospacers into a CRISPR-active and a CRISPR-inactive strain (Stringer *et al*., 2020a). Comparing the efficiency of conjugation for each protospacer into the two strains allows for quantification of the inhibitory effect of the CRISPR-Cas system; actively targeted plasmids conjugate efficiently into the CRISPR-inactive strain but inefficiently into the CRISPR-active strain. Our previous data also showed that there is considerable variability in the effectiveness of the different spacers in the *V. cholerae* A50 CRISPR array. We reasoned that a similar approach could be used to compare the effectiveness of individual *V. cholerae* A50 spacers against 64 plasmid-borne protospacers that are identical except for the trinucleotide sequence in the PAM position.

We constructed two plasmid libraries. Each library comprised 64 plasmids, one plasmid with each of the possible trinucleotide sequences in the PAM position. One library had a protospacer matching spacer #4 of the *V. cholerae* A50 CRISPR array; the other had a protospacer matching spacer #21. We previously showed that spacer #4 is ∼10-fold more effective than spacer #21 in preventing conjugation of a plasmid with a cognate protospacer (Stringer *et al*., 2020a), and we reasoned that this difference may be useful in distinguishing between the effectiveness of different PAM sequences. We pooled the two libraries together with a control plasmid that lacks a protospacer, and conjugated the plasmid pool into a CRISPR-inactive strain (*V. cholerae KS808* Δ*cas3-cas2*::*kan*^*R*^) and a CRISPR-active strain (*V. cholerae KS808* Δ*cas1*::*kan*^*R*^); both strains lack *cas1*, to prevent adaptation. We then quantified the abundance of each plasmid in the CRISPR-inactive and CRISPR-active strains by PCR-amplifying and sequencing the sequence surrounding and including the protospacer. We quantified the conjugation efficiency of each protospacer-containing plasmid by first comparing the number of sequence reads obtained from the CRISPR-inactive and CRISPR-active strains, and then normalizing to the control plasmid that lacks a protospacer and hence should be unaffected by the CRISPR-Cas system (Tables S1 and S2). Figure 1 shows the normalized conjugation efficiency associated with all possible trinucleotide sequences in the PAM position for the protospacers matching spacer #4 (*x*-axis) and spacer #21 (*y*-axis).

**Figure 1.**
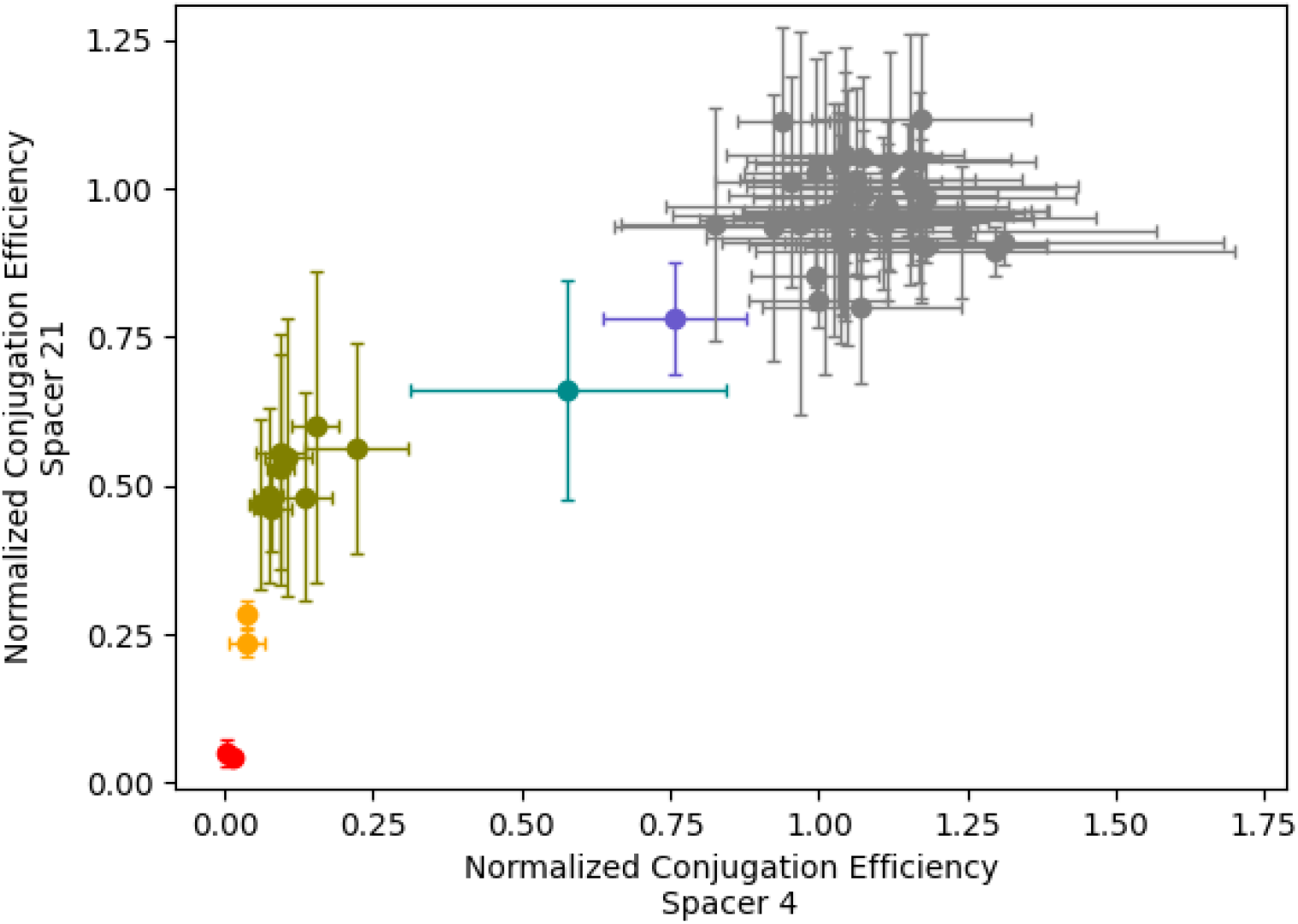
Conjugation efficiency for protospacer plasmids with each of the 64 possible NNN sequences in the PAM position. Two plasmid libraries were constructed: one with a protospacer matching the *V. cholerae* A50 spacer #4, and the other with a protospacer matching #21. Each of the two libraries contained all possible NNN sequences in the PAM position, i.e., 64 plasmids in each library. The scatter-plot shows the normalized efficiency of conjugation of each plasmid into a CRISPR-active strain of *V. cholerae*. Each datapoint represents a different NNN sequence in the PAM position. Values on the *x*-axis indicate conjugation efficiencies for plasmids with protospacers matching spacer #4. Values on the *y*-axis indicate conjugation efficiencies for plasmids with protospacers matching spacer #21. Datapoints for functional PAMs are grouped by color, as follows: AAC, AAT; ATT, ATC; TAC, CAT, CAC, GAT, TAT, GAC, AGC, AAG, AGT; AAA; AGG. Datapoints for all other NNN sequences (i.e., non-functional PAMs) are shown in gray. Values plotted are the mean of two independent biological replicates. Error bars indicate ± one standard deviation from the mean.

Overall, conjugation efficiencies for protospacers with active PAMs were up to 12-fold lower for the protospacer matching spacer #4 than the protospacer matching spacer #21, consistent with our earlier study (Stringer *et al*., 2020a). After accounting for the overall difference in CRISPR activity from spacer #4 versus spacer #12, the hierarchy of PAM efficacy was similar for the two protospacers. Our data confirm that AAY is the optimal PAM: AAC and AAT had the lowest conjugation efficiencies for both protospacers. 50 trinucleotide sequences were associated with conjugation efficiencies similar to those of the control plasmid that lacks a protospacer, strongly suggesting that these sequences cannot function as PAMs. 12 sequences had intermediate conjugation efficiencies, indicating that they function as suboptimal PAMs. ATY sequences were the strongest suboptimal PAMs, with conjugation efficiencies ∼5-8-fold lower than those for protospacers with AAY PAMs. Next were the NAY, AGY, and AAG PAMs, with efficiencies ∼9-50-fold lower than those for protospacers with AAY PAMs. The AAA PAM had efficiencies 13-130-fold lower than those for protospacers with AAY PAMs, and the AGG PAM had efficiencies 15-170-fold lower than those for protospacers with AAY PAMs.

Consider the three positions within the PAM: -3, -2, -1, with numbers indicating position relative to the start of the crRNA-DNA hybrid. Our data suggest that:

- A is the preferred base at position -3, but all bases are tolerated, with little difference between C/G/T.
- A is the preferred base at position -2, T is well-tolerated, G is weakly tolerated, but C renders the PAM non-functional.
- Y is the strongly preferred base at position -1, G/A are weakly tolerated, but C renders the PAM non-functional.

Based on the hierarchy of PAMs identified in this study, it should be possible to predict whether putative protospacers will be effectively targeted by the *V. cholerae* CRISPR-Cas system. Moreover, our work may facilitate the design protospacers with PAMs that function only weakly in interference but strongly promote primed adaptation. And our data may provide a framework for tuning the activity of technological applications that use the *V. cholerae* CRISPR-Cas system.

## MATERIALS AND METHODS

### Strains and Plasmids

All strains and plasmids used in this work are listed in Table 1. All oligonucleotides used in this work are listed in Table S3.

**Table 1.**
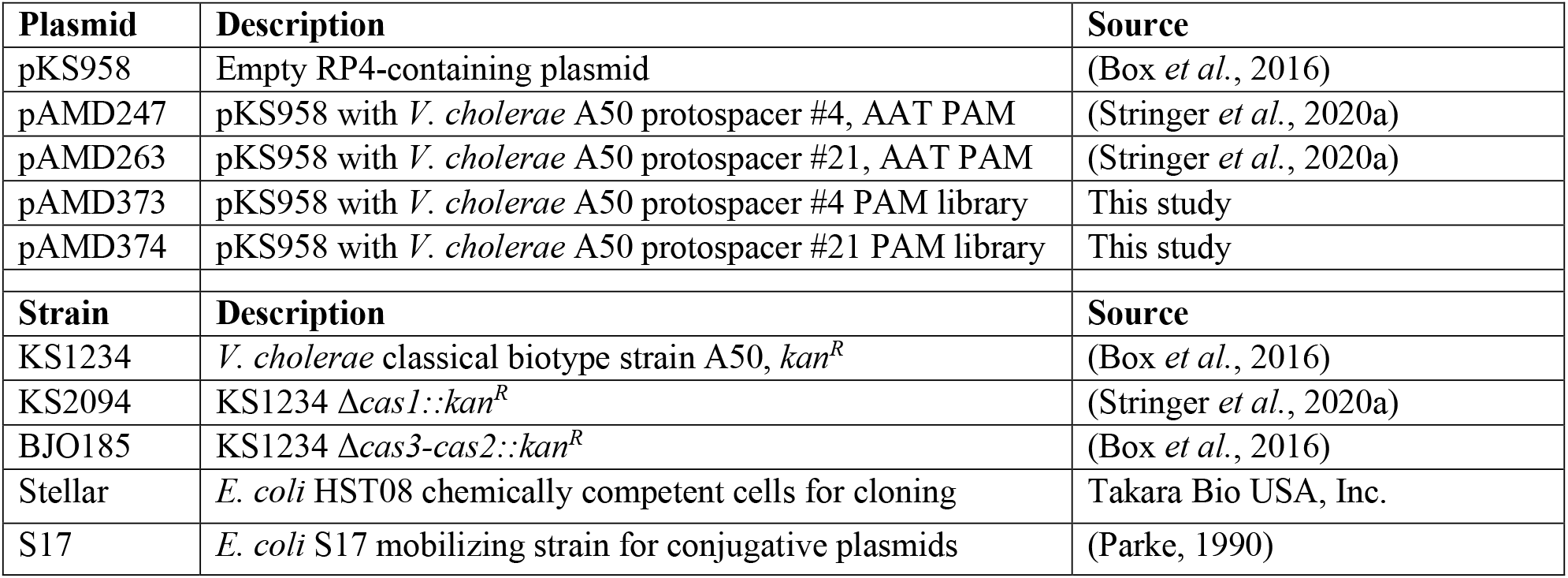
List of strains and plasmids used in this study.

### Cloning protospacer libraries

Protospacer #4 with a random trinucleotide sequence in the PAM position was PCR-amplified from pAMD247 using oligonucleotides JW10804 and JW10805. Protospacer #21 with a random trinucleotide sequence in the PAM position was PCR-amplified from pAMD263 using oligonucleotides JW10806 and JW10807. The PCR products were treated with *Dpn*I and cloned into *Hind*III/*Sph*I-digested pKS958 using an In-Fusion HD cloning kit (Takara Bio USA, Inc.), transforming into chemically competent *E. coli* Stellar cells plated on LB agar supplemented with 30 ug/mL chloramphenicol. Colonies were scraped, and plasmid was purified and transformed into *E. coli* S17.

### Conjugation of plasmids into *V. cholerae* and sequencing library construction

*E. coli* S17 cultures containing each of the protospacer libraries were grown separately in LB supplemented with 30 μg/mL chloramphenicol to an OD_600_ of 1.0, then pooled equally. The plasmid libraries were transferred from *E. coli* S17 to *V. cholerae* strains KS2094 and BJO185 by conjugation, as previously described (Box *et al*., 2016). 100 μL of cells from each conjugation were plated on LB agar supplemented with 75 μg/mL kanamycin and 2.5 μg/mL chloramphenicol. The next day, cells were scraped, resuspended in LB, and pelleted. ∼1 μL from each pellet was resuspended in 20 μL dH_2_O and incubated at 95 °C for 10 minutes. From each cell resuspension, 1.0 μL was used as a template in a PCR reaction with universal forward Illumina oligonucleotide JW10344 and reverse Illumina index oligonucleotides JW10345, JW10346, JW10347, or JW10348. Sequencing was performed using an Illumina MiSeq instrument (Wadsworth Center Applied Genomic Technologies Core). Two biological replicates were performed for all experiments.

### Analysis of sequencing datasets

Illumina sequencing .fastq files were analyzed using custom Python code (available from https://github.com/wade-lab/Vibrio-cholerae-PAM). Briefly, for each dataset we determined the number of sequence reads matching each possible trinucleotide sequence in the PAM position. We then normalized sequence read counts in the CRISPR-active strain to the corresponding values in the CRISPR-inactive strain. Lastly, we calculated normalized conjugation efficiencies for each trinucleotide sequence in the PAM position by dividing values for each sequence by those for the empty vector control (note that all plasmid libraries included empty vector as a byproduct of the cloning procedure).

## DATA ACCESSIBILITY

Raw Illumina sequencing data are available from the EBI ArrayExpress repository using accession number E-MTAB-14044.

## ACKNOWLEDGEMENTS

This study was supported by NIH grants R01GM122836 and R35GM144328 to J.T.W. We thank Kimberley Seed for helpful discussions and technical advice. We thank the Wadsworth Center Applied Genomic Technologies Core Facility and the Wadsworth Center Media and Tissue Culture Core Facility.

## SUPPLEMENTARY TABLES

**Table S1. Raw sequence read counts for protospacer plasmids**.

**Table S2. Normalized conjugation efficiencies for protospacer plasmids with each of the 64 possible NNN sequences in the PAM position**.

**Table S3. Oligonucleotides used in this study**.

